# An essential RNA-binding lysine residue in the Nab3 RRM domain undergoes mono and trimethylation

**DOI:** 10.1101/592923

**Authors:** Kwan Yin Lee, Anand Chopra, Kyle Biggar, Marc D. Meneghini

## Abstract

The Nrd1-Nab3-Sen1 (NNS) complex integrates molecular inputs to direct termination of noncoding transcription in budding yeast. NNS is positively regulated by methylation of histone H3 lysine-4 as well as through Nrd1 binding to the initiating form of RNA PolII. These cues collaborate with Nrd1 and Nab3 binding to target RNA sequences in nascent transcripts through their RRM RNA recognition motifs. In this study, we identify nine lysine residues distributed amongst Nrd1, Nab3, and Sen1 that are mono-, di-, or trimethylated, suggesting novel molecular inputs for NNS regulation. One of these methylated residues, Nab3 lysine-363 (K363), resides within its RRM, and is known to physically contact target RNA. Although mutation of Nab3-K363 to arginine (Nab3-K363R) causes a severe growth defect, it nevertheless produces a stable protein that is incorporated into the NNS complex, suggesting that RNA binding through Nab3-K363 is crucial for NNS function. Consistent with this hypothesis, K363R mutation decreases RNA binding of the Nab3 RRM *in vitro* and causes transcription termination defects *in vivo.* These findings reveal crucial roles for Nab3-K363 and suggest that methylation of this residue may modulate NNS activity through its impact on Nab3 RNA binding.

## INTRODUCTION

RNA PolII transcriptional termination is controlled through two distinct mechanisms in the budding yeast *Saccharomyces cerevisiae* (1-4). The first acts through the cleavage and polyadenylation factor/cleavage factor complex (CPF/CF), and couples the termination of protein encoding transcripts with their polyadenylation and nuclear export (5-7). The second mechanism functions through the Nrd1-Nab3-Sen1 (NNS) complex. NNS targets short noncoding RNAs such as snRNAs, snoRNAs, and cryptic unstable transcripts (CUTS) for transcriptional termination, following which the transcripts are targeted for processing or degradation through the Exosome complex (8-11). NNS has a major role in the control of pervasive non-coding RNA transcription, which if left unregulated can interfere with the transcription of protein coding genes (8,9,11-13).

Multiple regulatory mechanisms function to restrict CPF/CF and NNS termination to their respective target genes. CPF/CF terminates at 3’ ends of genes through its recognition of poly-A signals in emerging RNA transcripts (6,14-16). CPF/CF is further regulated through binding of its Pcf11 subunit to the serine-2 phosphorylated form of the PolII carboxy-terminal heptad repeat domain (CTD), which associates with transcriptional elongation (17-19). The action of CPF/CF cleaves the elongating RNA transcript, leading to the processing of these nascent mRNAs for polyadenylation (20). In accordance with its action on short noncoding transcripts, NNS termination is promoted through interaction of the Nrd1 CTD interaction domain (CID) with the serine-5 phosphorylated form of CTD, which is associated with the initiating/early-elongating form of RNA PolII (19,21,22). Moreover, genetic evidence suggests that NNS is positively regulated through methylation of histone H3 on lysine-4 (H3K4me), a chromatin mark widely associated with transcriptional initiation (23-30). By integrating these signals together with Nrd1 and Nab3 binding to cognate RNA sequences, NNS is thought to dislodge PolII from DNA in a manner that employs Sen1 ATPase activity (31-34).

As Nab3 and Nrd1 RNA recognition sites are found broadly in the transcriptome (35), and the additional molecular cues that promote NNS, -H3K4 methylation (23,24) and PolII CTD serine-5 phosphorylation-(21,22), are generic features of all PolII transcribed regions (36), additional mechanisms must act to restrict NNS from inappropriate termination. Here we illuminate a potential new mode of NNS regulation through lysine methylation of its subunits. Using LC-MS/MS full scan and SRM-MS mass-spectrometry approaches, we identify nine lysine residues distributed amongst Nrd1, Nab3, and Sen1 that exhibit methylated forms. Many of these lysine residues are found within conspicuous protein domains of regulatory potential. We focus this study on the Nab3-K363 methylation site, which resides within the Nab3 RRM and is known to make contact with the RNA backbone of target transcripts (37,38). *NAB3* is an essential gene (39), and we find that mutation of Nab3-K363 to alanine (Nab3-K363A) leads to the production of a stable protein but nevertheless causes lethality. Mutation of Nab3-K363 to its most structurally similar residue arginine (Nab3-K363R) results in viable cells of greatly reduced health. The slow growth of Nab3-K363R is associated with transcription termination defects *in vivo* and reduced RNA binding affinity *in vitro*. Our findings reveal that the integrity of Nab3-K363 is crucial for NNS function because of its role in Nab3 RNA binding. The mono and trimethylated forms of Nab3-K363 we have identified may thus have regulatory potential via an impact on Nab3 RRM RNA binding.

## MATERIALS AND METHODS

### Strains, media and plasmids

Standard *S. cerevisiae* genetic and strain manipulation techniques were used for strain construction and husbandry. Refer to Table S1 for strains and plasmids used in this paper. *NAB3* and *NRD1* containing 600bp of flanking sequences were cloned into pRS313 and pRS316. Cloning was performed using *in vivo* homologous recombination in yeast (40). Briefly, the parental plasmids were linearized with BamHI at the multiple cloning site and then transformed into budding yeast together with *NAB3* (600bp up and down) or *NRD1* (600bp up and down) containing 45bp of homology to the BamHI cut site sequences produced by PCR amplification. Transformants that grew on -HIS dropout media (for pRS313) and -URA dropout media (for pRS316) were screened by PCR and verified by sequencing. All plasmids were tested for their ability to complement their respective deletion mutants. The pRS313-*SEN1* and pRS316-*SEN1* plasmids were acquired from Dr. David A. Brow (University of Wisconsin) (41). Q5 site-directed mutagenesis (NEB) was used to introduce nucleotide changes that translate to single amino acid substitutions into pRS313 plasmids harboring *NAB3, NRD1*, and *SEN1*. For protein expression and purification from *E. coli*, Nab3-RRM encoding sequences corresponding to residues 329–419 were amplified by PCR using the primer Nab3_329_forward 5’-GCATCATATGAAGTCAAGATTATTCATTGG-3’ and Nab3_419_reverse 5’-GCGCGGCCGCTTAAGTAGAACTACTGTTTGTACC-3’ from genomic *cerevisiae* DNA (37). After restriction digestion using NdeI and NotI (NEB), PCR products were ligated into the pET28a expression vector DNA resulting in an N-terminal fusion with a hexahistidine tag. Q5 site-directed mutagenesis (NEB) was used to introduce the K363R, K363A, and S399A mutations into pET28a-NAB3-RRM(329-419). All plasmids were sequence verified.

### Serial dilution assay

Yeast strains were inoculated into several mL of -HIS -URA drop out media (YNB media (Multicell Wisent) containing 5 g/L of ammonium sulfate, -HIS -URA powder, 2% glucose) and grown overnight at 30°C. Each strain was diluted to an OD_600_ = 0.4, serially diluted five times, and spotted onto agar plates containing -HIS -URA dropout media 2% glucose and also on synthetic complete media 2% glucose supplemented with 0.1% 5-Fluoroorotic Acid (5FOA).

### RNA extraction and RT-qPCR analysis

Strains were grown to mid-logarithmic phase and 5-10 OD_600_ equivalents of cells were harvested for RNA extraction. RNA was extracted with acidic phenol at 65°C for 30 min. RNA was then purified, precipitated, and resuspended in RNase free water. cDNA was prepared using either random nonamers or site-specific primers and Maxima H Minus Reverse Transcriptase (ThermoFischer Scientific) according to the manufacturer’s instructions. PCR amplification of cDNA was detected using SYBR green in the BioRad iQ5 Multicolor Real Time PCR Detection System. All primer sequences are detailed in Table S2. qPCR signal was normalized to the reference transcript *ACT1*.

### Nab3-RRM purification

BL21(DE3) *E coli.* cells containing pET28a-NAB3-RRM(329-419) (or mutant plasmids) were grown in 750 mL LB + 50 µg/mL kanamycin to OD_600_ = 0.4-0.6. Protein expression from pET28a plasmids was induced with 1 mM IPTG overnight at 16°C. Cells were then harvested into two 375 mL pellets stored in −80°C. Nab3-RRM peptides were purified using Nickel Column Purification. Cell pellets were resuspended in PBS with protease inhibitors (E64, Bestatin, Pepstatin A, and PMSF), 1 mg/mL Lysozyme, 2% Triton X-100, and RNase free DNaseI. Lysis was achieved by first sonicating for 4 cycles 30s ON/OFF and then by incubating the cells for 20 min on a rotator at 37°C. Cell debris was clarified by centrifugation at 38720xg for 30 min. Cleared lysates were then passed through a Ni-NTA column (Qiagen 30250) that was equilibrated with P5 Buffer pH 7 (20 mM NaHPO_4_, 500 mM NaCl, 10% Glycerol, 0.05% Triton X-100, 5 mM Imidazole, and 1 mM DTT). The column was then washed three times with P40 Wash Buffer pH 7 (20 mM NaHPO_4_, 500 mM NaCl, 10% Glycerol, 0.05% Triton X-100, 40 mM Imidazole, and 1 mM DTT). Elution was accomplished by passing 5 mL of P250 Elution Buffer pH 7 (20 mM NaHPO_4_, 500 mM NaCl, 10% Glycerol, 0.05% Triton X-100, 250 mM Imidazole, and 1 mM DTT) through the Ni-NTA column. Purified protein was dialyzed into buffer containing 20 mM Tris-HCl pH 8, 200 mM NaCl, 10% Glycerol, 1 mM DTT and used for downstream applications. Protein concentration was determined by a standard Bradford assay.

### RNA binding assay

Purified 6xHis-tagged Nab3 329-419 WT, K363R, K363A, and S399A proteins were incubated with 25 µM biotinylated RNA probe (snR47; 5’-UUUCUUUUUUCUUAUUCUUAUU-3’) in binding buffer (10 mM Tris pH 7.8, 50 mM NaCl, 1 mM EDTA, 5% Glycerol, 2.5 mM DTT) for 30 min at 15°C. Streptavidin-agarose beads (Novagen), blocked with bovine serum albumin, were added and rotated for 2h at 4°C. Beads were washed four times with binding buffer containing 0.1% Tween-20 and bound proteins were eluted with 2X Laemmli Sample Buffer (Bio-Rad Laboratories). Eluted proteins were subjected to electrophoresis on a 17% resolving gel at 120V for 2 hrs and transferred to PVDF membrane at 180mA for 2 hrs and immunoblotted with His-Probe-HRP (Pierce) at a 1:5000 dilution. Immunoreactive bands were detected by chemiluminescence on a BioRad Chemidoc MP imager and quantified based on relative densitometry. The equilibrium dissociation constant (K_d_) was determined based on densitometry of the visualized gel shift as previously published (42-44).

### Protein Extraction and Immunoblot Analysis

One mL of logarithmically growing yeast cells was harvested for protein extraction. Cells were immediately pelleted and resuspended into 250 μL of 0.1 N NaOH for 5 min. The NaOH was then removed by centrifugation so that the cell pellet could be resuspended in 1 x Laemmli Sample Buffer. This resuspension was boiled for 5 min. Total protein concentration was determined using an RC/DC assay (BioRad 5000121). Equal amounts of protein were electrophoresed on 8% or 10% SDS-PAGE gels and transferred onto Immobilon-PVDF Transfer membranes. (Millipore IPVH00010). Immunoblot analysis was performed using standard procedures. All blots were scanned with a ChemiDoc XRS+ Imaging System. Band intensities were quantified using ImageJ 1.51 v software.

Sen1 immunoblots were performed with slight differences from the standard protocol above as described previously (41). First, samples were electrophoresed using 4–15% Mini-PROTEAN TGX Precast Gels (Bio-Rad 4561086) at 140 V for 1 hr transferred onto Immobilon-PVDF Transfer membranes. Blots were blocked with 5% dried milk in TBST buffer with 0.1% Tween-20 at 23°C for 1 hr, incubated with Sen1 antibody (1:2000 dilution) for 1 hr and then an anti-rabbit secondary at 1:3000. Blots were scanned with a ChemiDoc XRS+ Imaging System and band intensities were quantified using ImageJ 1.51 v software.

### Immunoprecipitation

Co-immunoprecipitation was performed as described previously (45). Cell pellets containing 0.5 g of cells were collected from yeast cells grown to OD_6oo_ = 1-2. Pellet were frozen at −80°C until they were ready to be used. Frozen cell pellets were resuspended in Lysis Buffer (50 mM Na-HEPES, pH 7.5, 200 mM NaOAc, pH 7.5, 1 mM EDTA, 1 mM EGTA, 5 mM MgOAc, 5% glycerol, 0.25% NP-40, 3 mM DTT, 1 mM PMSF, and protease inhibitor cocktail (Roche 11836170001)) and lysed by three rounds of bead beating (BioSpec Mini-beadbeater-16), 1 min on and 1 min off on ice. Cell lysates were clarified by centrifugation. Total protein in the whole cell lysates was quantified using the RC/DC assay (BioRad 5000121). All samples were adjusted to the same concentration by adding Lysis Buffer. 30 μL of Protein A-Sepharose (15 μL of packed beads in a 50% slurry) was used per Anti-Nrd1 IP. 30 μL of Protein G-Sepharose (15 μL of packed beads in a 50% slurry) was used per Anti-Nab3 IP. Protein A-Sepharose beads were washed several times with Lysis Buffer and finally resuspended to 200 μL with 10 μL of anti-Nrd1 antibody. Protein G-Sepharose was put through the same protocol but with 5μL of anti-Nab3 antibody. 500 μL of whole cell lysate was then added to the 200 μL of beads and antibody. Samples were placed on an end-over-end rotator at 4°C for 2 hr. Beads were gently collected by low speed centrifugation and wash several times using Wash Buffer (50 mM Na-HEPES, pH 7.5, 200 mM NaOAc, pH 7.5, 1 mM EDTA, 1 mM EGTA, 5 mM MgOAc, 5% glycerol, 0.25% NP-40, 3 mM DTT, and 1 mM PMSF). Samples were eluted by suspending beads in 50 μL 1 x Laemmli Sample Buffer followed by a 65°C incubation for 10 min. Eluted proteins were immunoblotted for Nrd1, Nab3, Sen1, and Pgk1 as described above.

### Antibodies

An Anti-Nab3 (2F12) monoclonal mouse antibody was acquired from Dr. Maurice Swanson at the University of Florida and described previously (39). Anti-Nrd1 and anti-Sen1 rabbit antibodies were acquired from Dr. David A. Brow at the University of Wisconsin (41). The Anti-Pgk1 antibody (ab113687) was purchased from Abcam. Antibodies were used at the following dilutions for immunoblot analysis: Nab3 (1:5000), Nrd1 (1:5000), Sen1 (1:5000), and Pgk1 (1:5000).

### TAP purification

Two step TAP purification was used to isolate Nrd1, Nab3, and Sen1 as previously described (46). Exponential cultures were grown overnight to OD_600_ between 1-2. Cells were harvested by centrifugation then washed once with cold water, once with cold Yeast Extract Buffer (YEB) (100 mM HEPES-KOH pH 7.9, 240 mM KCl, 5 mM EDTA, 5 mM (EGTA)-KOH pH 7.9, and 2.5 mM DTT), and once with cold YEB buffer containing protease inhibitor. For future use, cells were snap frozen in Falcon Tubes using liquid nitrogen. Cell lysis was performed by mechanically grinding the cell pellets into a fine powder with dry ice. The lysed powder was resuspended into an equal volume of YEB buffer containing protease inhibitor. Cell lysates were subject to ultracentrifugation to remove debris followed by dialysis to a buffer suitable for IgG binding (Dialysis buffer: 10 mM Tris–HCl pH 7.9, 100 mM NaCl, 0.2 mM EDTA, 0.5 mM DTT, and 20% glycerol). TAP-tagged proteins were bound to IgG Sepharose beads that recognize the protein A moiety on the TAP tag. Protein associated with the IgG Sepharose beads were eluted using TEV protease, an enzyme that cleaves a linker region on the TAP tag, removing the protein A moiety permanently. In the second purification step, proteins that were freed from IgG were bound to Calmodulin Sepharose in a calcium dependent manner and subsequently eluted using EGTA. SDS-PAGE and silver staining were used to verify the presence of isolated protein complexes.

### Mass Spectrometry

Samples were first digested using trypsin (Roche Diagnostics) overnight at 37 °C. Digests were desalted and dried in a SpeedVac. After desalting on C18-Zip Tip and drying, samples were loaded onto a Thermo Easy-Spray analytical column (75 µm i.d. × 500 mm) C18 column with an Easy-nLC 1000 chromatography pump. For each analysis, we reconstituted peptides in 20 µL of 0.1% FA and loaded 4 µL onto the column. Peptides were separated on a 125 min (5–40% acetonitrile) gradient. Mass spectra were collected on a Q-Exactive hybrid quadrupole-Orbitrap mass spectrometer coupled to an Easy-nLC 1000 system (ThermoFisher). The spectrometer was set in full MS/data-dependent-MS2 TopN mode: mass analyzer over *m*/*z* range of 400–1600 with a mass resolution of 70□000 (at *m*/*z* = 200), 35 NEC (normalized collision energy), 2.0 *m*/*z* isolation window, and 15 sec dynamic exclusion. The PeaksX software (Bioinformatics Solutions Inc.) was used for data processing.

For SRM MS, the *in silico* protease digest patterns and the corresponding MRM transitions were compiled with the Skyline software (47). Transitions that are larger than the precursor ion were selected on the basis of the Skyline predictions and the specific ions that allow unambiguous identification of the methylated lysine site were included. An isolation list was exported and used for the SRM MS method. Samples for SRM MS were digested and treated as mentioned above. The Skyline software was used for data processing.

## RESULTS

### Identification and characterization of NNS methylated lysine residues

We previously showed that NNS function was negatively regulated through one or more histone lysine methyltransferases in a manner independent of histone methylation (24). To investigate this, we used a proteomic approach to identify components of the NNS complex that undergo lysine methylation. NNS was purified by tandem affinity purification using Nab3-TAP (Figure 1A). The purified NNS complex was subjected to LC-MS/MS and we identified nine lysines distributed amongst the three subunits that were mono-, di-, or trimethylated (Figure 1B, 1C, and S1). Many of these modified lysine residues map to functional domains of NNS subunits (Figure 1D). For example, Nab3-K213me1 maps within a domain of Nab3 that mediates its interaction with Nrd1 (Figure 1D) (10). Moreover, both K363me1 and K393me2 of Nab3 reside within its RRM (Figure 1D) (48). Lysine methylations on Nrd1 also map to functional domains (Figure 1D). Nrd1-K148me3 is located within its CID, while Nrd1-K171me1 maps to its Nab3 interaction domain (Figure 1D) (10). Three methylated lysine residues were identified within Sen1 (Figure 1C and 1D). Interestingly, both Sen1-K19 and Sen1-K21 exhibit both mono and dimethylated forms (Figure 1C and 1D). Although the N-terminus of Sen1 possesses no predicted secondary structure, two-hybrid analysis identified physical interactions of this region with the Rad2 and Rnt1 proteins, which are involved in DNA repair and RNA processing (49). Sen1-K1921me1 resides within one of Sen1’s two functionally confirmed nuclear localization sequences (Figure 1D) (41,50).

**Figure 1.**
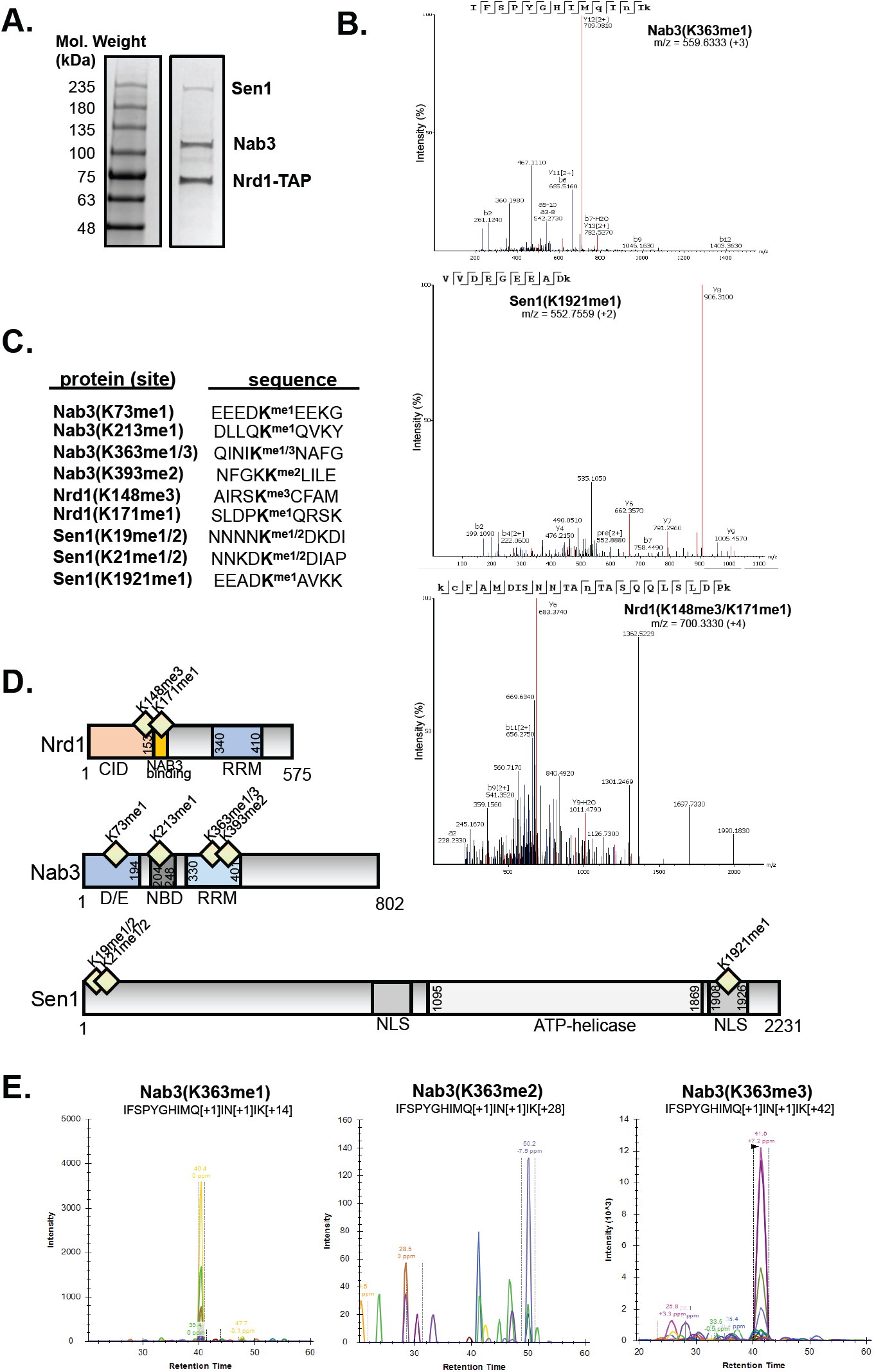
Multiple lysine residues on Nrd1, Nab3, and Sen1 are methylated. (A) The Nab3-Nrd1-Sen1 (NNS) complex was affinity purified through Nab3 tandem affinity purification. A Coomassie stained SDS-PAGE gel is shown. (B) Representative mass spectra for the identification of novel Nrd1, Nab3, and Sen1 lysine methylations. Mass spectra for Nab3(K363me1), Nrd1(K148me1&K171me1) tandem methylations, and Sen1(K1921me1) are shown. (C) A total of nine NNS methylation events were discovered by LC-MS/MS (QE Orbitrap) and post-analysis using PEAKS Studio software. (D) Nrd1, Nab3, and Sen1 lysine methylations mapped to functional domains of their respective proteins. (E) Methylation status of Nab3(K363) exists as a mono and trimethyl modification. Chromatographs show representative transition ions detected by SRM-MS. Abbreviations; CID (CTD-interacting domain), D/E (D/E-rich), NBD (Nrd1 binding domain), RRM (RNA recognition motif), NLS (nuclear localization sequence).

To assess whether any of the lysine residues that undergo methylation exhibited a detectable role in NNS function, we mutated each residue to arginine (Figure S2). Substitution of lysine to arginine (K→R) is frequently employed in an attempt to mimic a constitutively unmodified lysine residue. As the components of the NNS complex are all encoded by essential genes, mutant alleles of NNS were introduced into yeast strains using a plasmid shuffling approach (51). In this approach, the viability of *nrd1Δ, nab3Δ*, and *sen1Δ* strains depended on the presence of their respective wildtype genes on low-copy *URA3*-based vectors. Mutant alleles of *nrd1, nab3*, and *sen1* were generated on a second low-copy *HIS3*-based vector and introduced into their respective deletion mutants. Thus, strains harboring both the *URA3* plasmid (wildtype) and the *HIS3* plasmid (mutant allele) were obtained. These strains were then spotted onto agar plates with -HIS -URA dropout media as a control and synthetic complete media containing 0.1% 5FOA to select against cells with the *URA3*-based vector. Cells grown on -HIS -URA dropout media maintain both vectors and all displayed comparable growth regardless of the contents of their *HIS3*-based vectors (Figure S2). As expected, the *HIS3-*based empty vector did not support growth, but *nrd1Δ, nab3Δ,* or *sen1Δ* were complemented with the *HIS3-*based vector containing their respective wildtype genes (Figure S2). No apparent growth defects were observed when Nrd1 and Sen1 lysine methylation sites were mutated to arginine (Figure S2A and S2C). For Nab3, we observed a severe growth defect when lysine-363 was mutated to arginine (Nab3-K363R), but no apparent phenotype for the Nab3-K73R, K213R, or K393R mutants (Figure 2A and S2). The Nab3-K363R phenotype similarly manifested when all four methylated lysines were mutated to arginine (Figure S2B). Western blotting revealed that all lysine to arginine mutant proteins were expressed stably (Figure 2B and S3).

**Figure 2.**
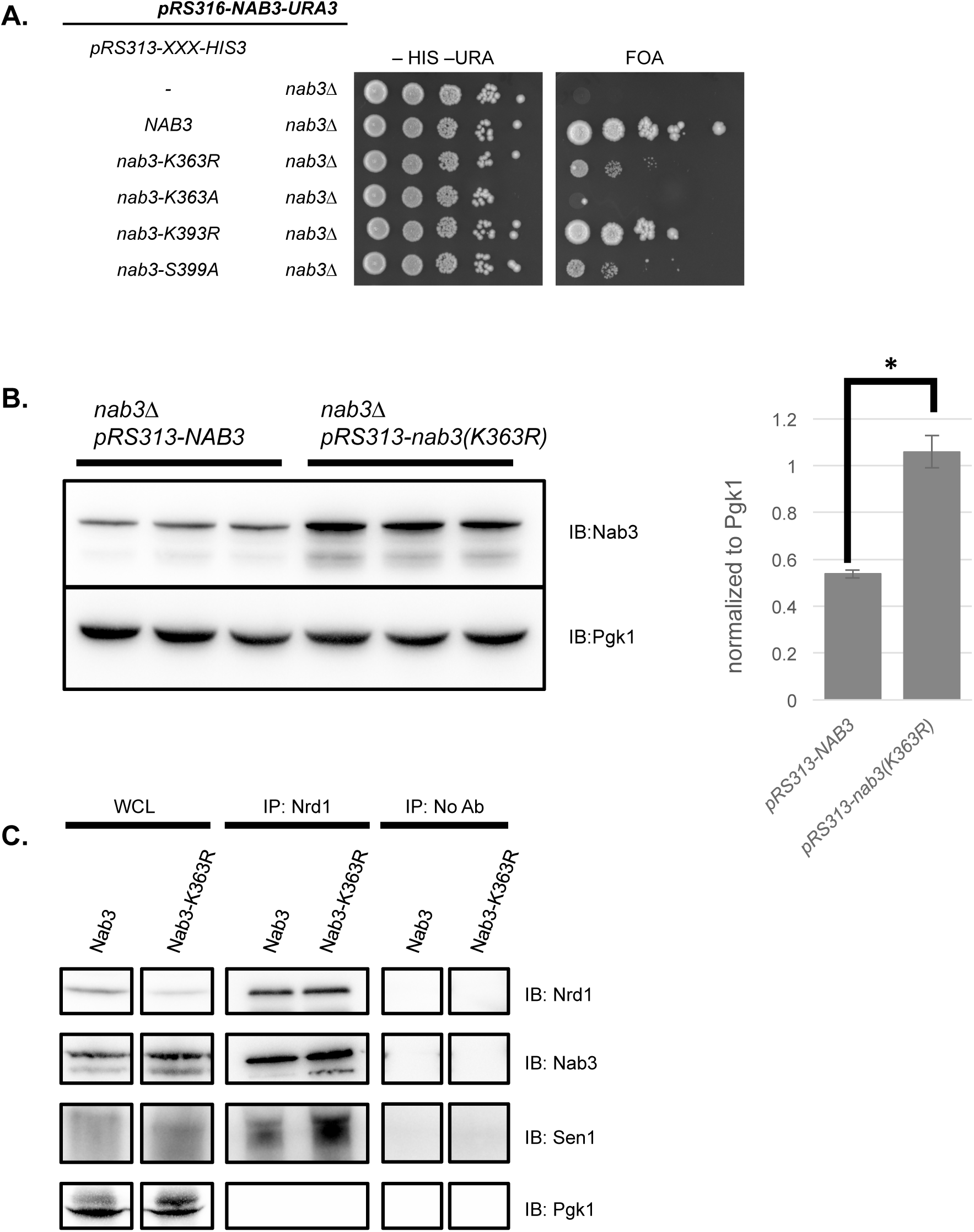
The Nab3 lysine 363 residue is important for cell viability. (A) Mutations in *NAB3* were introduced using a plasmid shuffling approach, in which a plasmid expressing a *NAB3* allele (pRS313-*nab3-allele-HIS3)* was transformed into a *nab3Δ* strain complemented with the pRS316-*NAB3-URA3* plasmid. Strains containing both plasmids were serially diluted ten-fold, spotted onto agar plates containing -HIS -URA dropout media and also on synthetic complete media supplemented with 5FOA to select against the pRS316-*NAB3-URA3* plasmid. From top to bottom: *pRS313-HIS3* (empty vector), *pRS313-NAB3-HIS3* (wildtype), *pRS313-nab3-K363R-HIS3, pRS313-nab3-K363A-HIS3, pRS313-nab3-K393R-HIS3,* and *pRS313-nab3-S399A-HIS*. K→R is a lysine to arginine substitution, K→A is a lysine to alanine substitution, S→A is a serine to alanine substitution. (B) Immunoblot analysis of Nab3 protein levels in wildtype and *nab3-K363R* mutants. Three biological replicates of each genotype are shown. Protein levels were quantified relative to the loading control Pgk1. Error bars represent standard deviation of three biological replicates, and significance was calculated busingy a two-tailed student’s t-test and denoted by *p<0.01. (C) Nrd1 immunoprecipitation in wildtype and *nab3-K363R* mutants. Total cell lysates (WCL) were immunoprecipitated with a polyclonal Nrd1 antibody (IP: Nrd1) or no antibody (No Ab) as a negative control followed by immunoblot analysis with anti-Nrd1, anti-Nab3, anti-Sen1, and anti-Pgk1 antibodies.

To more sensitively interrogate the K→R mutants, we also tested if they affected the growth of temperature sensitive mutants of other complex subunits in *trans* (for instance a *nab3-K73R* mutant in a *nrd1-102* mutant or a *sen1-1* mutant). Sen1 K→R mutants did not affect the growth of *nab3-11* or *nrd1-102* mutants; Nab3 K→R mutants did not affect the growth of *nrd1-102* or *sen1-1* mutants; and Nrd1 K→R mutants did not affect the growth of *nab3-11* or *sen1-1* mutants (data not shown). The only other phenotype we observed was a subtle but highly reproducible suppression of the *nrd1-102* allele by a cis mutation of K171 to arginine (Figure S4). This suggests that Nrd1-K171, and perhaps methylation of Nrd1-K171, has some repressive consequence for Nrd1 function.

### Nab3-K363R encoded a stable protein that was assembled into NNS

Of the nine methylated lysine residues we identified, only mutation of Nab3-K363 resulted in an apparent growth phenotype (Figure 2A, S2B). This result illuminated a functional significance of Nab3-K363 for its *in vivo* function, and we choose to focus on this residue for continued studies. To fully characterize the methylation profile of the Nab3-K363 site, we used SRM-MS with an independently purified NNS complex to monitor its methylation status in a targeted MS manner (Figure 1E). Using this approach, we found that Nab3-K363 exhibited mono and tri-methylated forms. The *nab3-K363R* mutant is similar to the well-studied *nab3-11* mutant in that both contain mutations in the RRM (F371L and P374L mutations for *nab3-11*) that cause slow growth and even temperature sensitivity for *nab3-11* (52). To highlight that the *nab3-K363R* phenotype was not a generic consequence of mutating any RRM residue, we note that the RRM associated Nab3-K393R mutant displayed wildtype-comparable growth (Figure 2A, S2B).

An explanation for the Nab3-K363R growth defect could be that it caused destabilization of the essential Nab3 protein. To test this, we measured Nab3 protein levels using immunoblotting. Surprisingly, we found that Nab3-K363R proteins levels were approximately two-fold higher than the wildtype Nab3 (Figure 2B and S3). This increase in Nab3-K363R protein levels was not due to an increase in mRNA abundance as *nab3-K363R* and *NAB3* transcript levels were equivalent (Figure S5). Next, we asked if Nab3-K363R was incorporated into NNS, positing that improper formation of the NNS complex might cause the severe cell growth defect caused by *nab3-K363R*. To test the integrity of the NNS complex in wildtype and *nab3-K363R* mutants, we used a polyclonal antibody to immunoprecipitate Nrd1. Immunoblotting was then used to assess the levels of associated Nab3 and Sen1 in these immunoprecipitates. We found that Nrd1 associated with Sen1 and Nab3 indistinguishably in WT and *nab3-K363R* strains (Figure 2C). A reciprocal immunoprecipitation was performed using a monoclonal Nab3 antibody, validating the interaction between Nrd1 with both wildtype Nab3 and Nab3-K363R (Figure S6).

These findings suggest that the integrity of Nab3-K363 was of crucial importance for Nab3 function, and that was not due to any consequences on NNS assembly. The K→R substitution was chosen as a conservative change that might mimic a constitutively unmodified lysine using the rationalization that both lysine and arginine are positively charged amino acids with sidechains that have similar structures. To further investigate if the chemical properties of Nab3-K363 were important for its function, we constructed a Nab3-K363 to alanine (K→A) mutant. The sidechains of lysine and alanine are completely different both in structure and chemical properties. Although *nab3-K363A* encoded a stable protein, a *nab3Δ* strain complemented with a plasmid expressing *nab3-K363A* was inviable (Figure 2A and S7). Collectively, these findings show that the precise biochemical characteristics of Nab3-K363 were crucial for proper NNS function, and that this was unrelated to protein stability or complex formation.

### Nab3-K363R caused transcription read-through defects

Our findings implied that, despite permitting the assembly of an NNS complex, Nab3-K363R caused reduced NNS function. We evaluated this by investigating termination defects caused by Nab3-K363R. The *snR13-TRS31* locus is commonly investigated for evidence of transcription read-through defects caused by reduced NNS function. *SNR13* encodes one of many snoRNA transcripts that are normally terminated by NNS (9,53). In NNS mutants, failed termination at *snR13* leads to a read-through transcript through the downstream *TRS31* gene, which is terminated by CPF/CF resulting in a concatenated *snR13-TRS31* transcript (9). To investigate whether *nab3-K363R* mutants caused defects in transcription termination, we measured transcription read-through at *snR13-TRS31* using a RT-qPCR approach. Initially, we used a *TRS31* gene specific primer to create cDNA from RNA isolated from wildtype, *nab3-K363R*, and a *nab3-11* control. With this strategy, wildtype cells should contain a cDNA corresponding to the *TRS31* mRNA and no *SNR13-TRS31* concatenated transcripts (Figure 3A). In contrast NNS mutants should contain significantly more *SNR13-TRS31* transcripts (Figure 3A). The amount of *SNR13-TRS31* cDNA was determined with quantitative polymerase chain reaction (qPCR) using a forward primer that anneals within *SNR13* and a reverse primer that anneals in the intergenic region between *SNR13* and *TRS31* (Figure 3A). The level of read-through transcripts in each strain was normalized to *ACT1* mRNA levels. In agreement with previous studies, we found that strains with the *nab3-11* temperature sensitive allele accumulated significant levels of read-through transcripts at *SNR13-TRS31*. Strains with the *nab3-11* allele accumulated 15-fold more read-through transcripts compared to wildtype (Figure 3A). In comparison, strains with the *nab3-K363R* allele accumulated three-fold more read-through transcripts than wildtype (Figure 3A).

**Figure 3.**
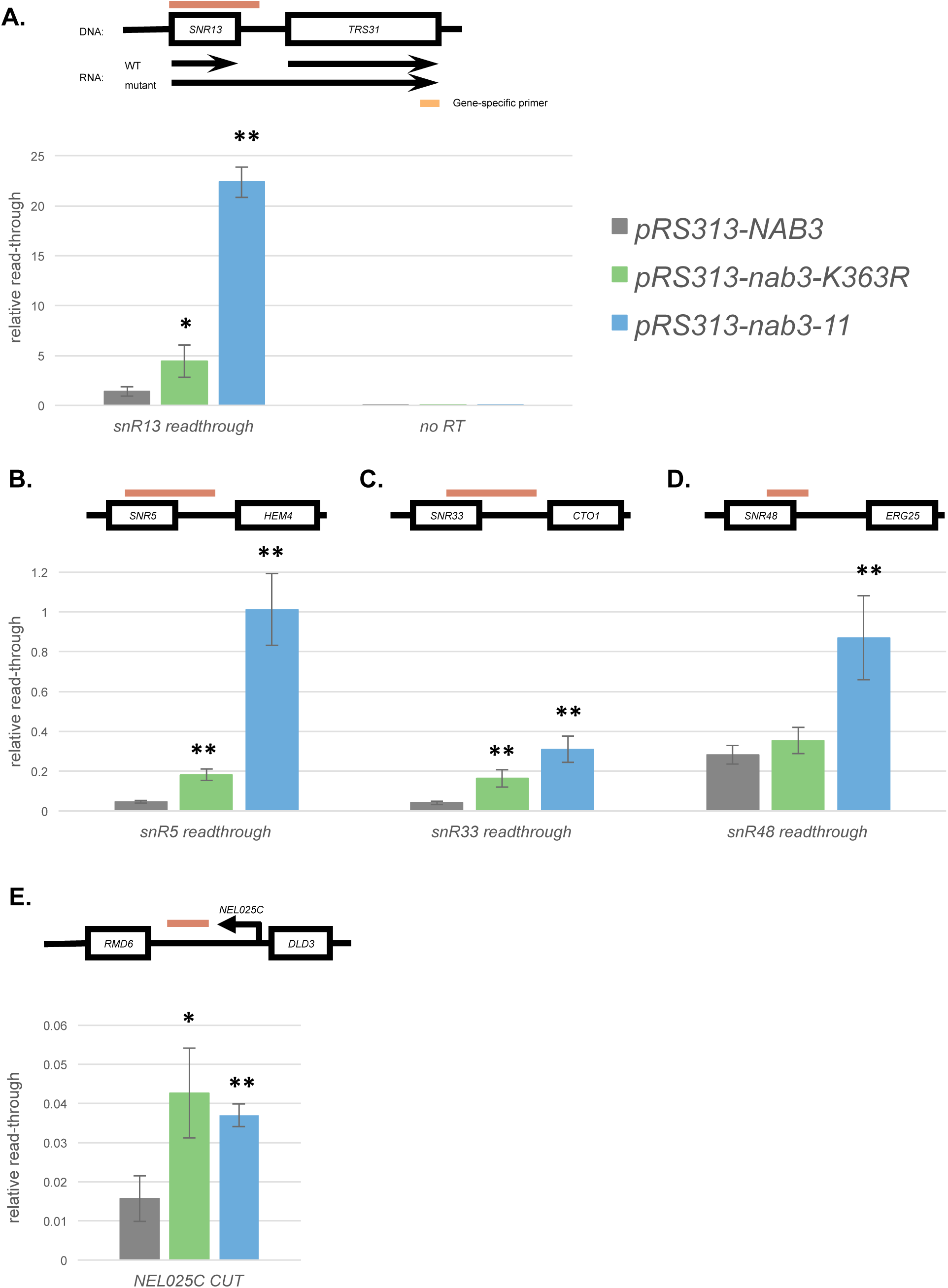
The Nab3-K363R mutant exhibits transcription termination defects at several noncoding RNAs. (A) Total RNA from *NAB3, nab3-K363R*, and *nab3-11* cells was processed into cDNA using a *TRS31* gene specific primer (illustrated as an orange rectangle). The presence of *SNR13* read-through transcription was analyzed by qPCR using a forward primer that anneals within the snoRNA and a reverse primer that anneals at a downstream intergenic region. The qPCR product is represented by the red rectangle. (B-E) Total RNA from *NAB3, nab3-K363R,* and *nab3-11* cells was processed into cDNA using random nonamers. Read-through transcription was measured at the following snoRNA genes: (B) *SNR5*, (C) *SNR33,* (D) *SNR48*. RT-qPCR was also used to measure levels of the (E) *NEL025C CUT*. qPCR signal from the read-through and *NEL025C* primers was normalized to *ACT1* mRNA levels, which appear comparable in all three strains. Error bars represent standard deviation of three biological replicates. Significance between each mutant and the WT control (*pRS313-NAB3)* is calculated by a two-tailed student’s t-test and denoted by * p<0.05, ** p<0.01. No reverse transcription control (No RT, n=1).

To assess accumulation of read-through transcripts at a wide range of loci, we subsequently tested a random primer approach to create cDNA. We measured the levels of *SNR13-TRS31* cDNA generated by random priming using the same qPCR primers as above. Data from gene specific priming and random priming were comparable, with a 10-fold and 2.5-fold more read-through transcript accumulation compared to wildtype in the *nab3-11* and *nab3-K363R* mutants respectively (Figure S8A). We similarly measured read-through transcript accumulation at a number of different snoRNA loci (Figure 3B, 3C, 3D, and S8B). Generally, read-through transcripts accumulate to higher levels in *nab3-11* strains compared with *nab3-K363R* strains. For instance, *snR5-HEM4* and *snR33-CTO1* loci follow the same trends as at *snR13-TRS31*, with a significant accumulation of read-through transcripts in *nab3-K363R* and *nab3-11* compared to wildtype (Figure 3B and 3C). Interestingly, at *snR47-YDR042C* and *snR48-ERG25*, we found that read-through transcript accumulation was elevated in *nab3-11* but not *nab3-K363R* (Figure 3D and S8B). We also examined the cryptic unstable transcript NEL025C, which is terminated by NNS, leading to its instability (11). In contrast to the snoRNA read-through transcripts, NEL025C CUT accumulated to similar levels in both the *nab3-11* and *nab3-K363R* mutants (Figure 3E). These results show that Nab3-K363R caused widespread NNS transcriptional termination defects.

### Nab3-K363R caused reduced RNA binding *in vitro*

The conspicuous location of Nab3-K363 within the RRM domain suggested that this residue might be important for RNA binding. Indeed, X-ray crystallographic and NMR spectroscopy studies of Nab3-RRM bound to its UCUU recognition sequence revealed that Nab3-K363 makes contact with the RNA backbone (37,38). Specifically, the sidechain of Nab3-K363 forms a hydrogen bond with one of the bridging phosphates of the RNA (Figure 4A) (37,38). These previous findings led us to propose that Nab3-K363R weakened the binding of Nab3 to its cognate RNA sequence. We tested this hypothesis using *in vitro* RNA pulldown assays, measuring the ability of bacterially expressed and purified wild-type and mutant Nab3 RRM proteins to bind to a biotinylated RNA probe corresponding to snR47. As expected, Nab3-RRM bound the probe robustly (Figure 4B, 4C, 4D). In agreement with a previous study, we did not detect RNA binding by a S399A RRM mutant protein, establishing the validity of our RNA binding assay (Figure 4B) (38). Intriguingly, although the Nab3 S399A mutant was previously reported to cause lethality in yeast (38), we were able to recover these mutants, though they were very slow growing (Figure 2A). Like with *nab3-K363A* and *nab3-K363R*, the *nab3-S399A* growth defect was not due to impaired protein expression (Figure S7).

**Figure 4.**
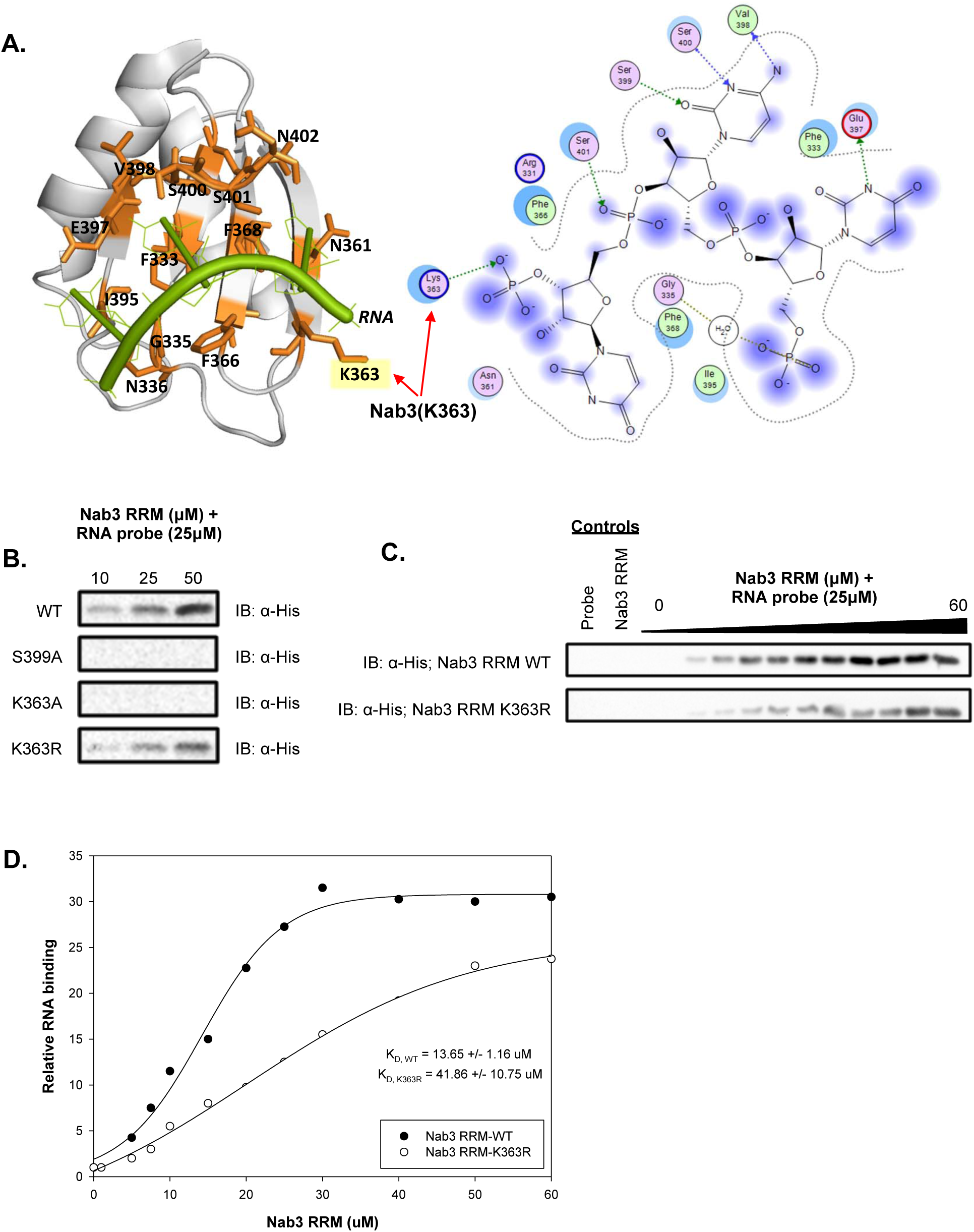
The Nab3 lysine-363 to arginine mutation attenuates RNA binding *in vitro*. (A) Co-crystal structure of Nab3 demonstrating the proximity (left) and interaction (right) of the K363 methylation site with bound RNA (PDB 2XNQ). Structures were visualized using PyMol (left) and Molecular Operating Environment (right). (B) Effect of single mutants (S399A, K363A, K363R) on RNA binding activity of Nab3 RRM (329-419) protein to 25μM snR47 RNA probe (UUUCUUUUUUCUUAUUCUUAUU). Immunoblot detection by terminal 6xHis tag present on all Nab3 constructs. (C) Dose-response (0-60μM) of Nab3 RRM (329-419) and Nab3 RRM (329-419; K363R) to 25μM snR47 RNA probe. (D) Quantification of Nab3 RRM binding activity to snR47 probe. All RNA binding was monitored by RNA pull-down assay.

Next, we assessed Nab3-K363A and -K363R RRM for their ability to bind snR47 RNA. Like with the S399A mutant, we found that the K363A RRM also exhibited no RNA binding activity *in vitro* (Figure 4B). In contrast, the K363R RRM maintained the ability the bind RNA (Figure 4B), but at a significantly decreased affinity. We quantified this decreased binding by calculating the equilibrium dissociation constants (K_d_) for the wildtype and K363R Nab3 RRMs. Nab3-RRM binds the snR47 RNA with moderate affinity with an apparent K_d_ of 13.65 μM (Figure 4C, 4D). The K363R mutation decreased snR47 RNA binding affinity of the Nab3 RRM by three-fold (K_d_ of 41.86 μM). The differences in viability of wildtype vs *nab3-K363A* and *nab3-K363R* strains thus mirror their respective RNA binding capacity, revealing that the contacts between K363 and the RNA backbone represent a highly sensitive interface controlling NNS function.

## DISCUSSION

Post-translational modifications (PTMs) are well known for their ability to impart protein regulation through a diverse array of molecular mechanisms. Although protein lysine methylation is most famously understood through studies of histone methylation, this modification has emerged as a PTM that impacts a wide array of proteins (54). Surprisingly given the known recruitment of numerous histone lysine methyltransferases to chromatin, the identification of methyllysines in chromatin-associated proteins is still nascent. We identify here nine methylated lysine residues in the NNS transcriptional termination complex and show that Nab3-K363 is of critical importance for NNS function through its role in mediating RNA binding. Although in need of further studies to confirm, our findings suggest a model in which mono and/or trimethylation of Nab3-K363 regulates NNS via impacting the ability of Nab3 to bind RNA (Figure 5). This will be discussed further below.

**Figure 5.**
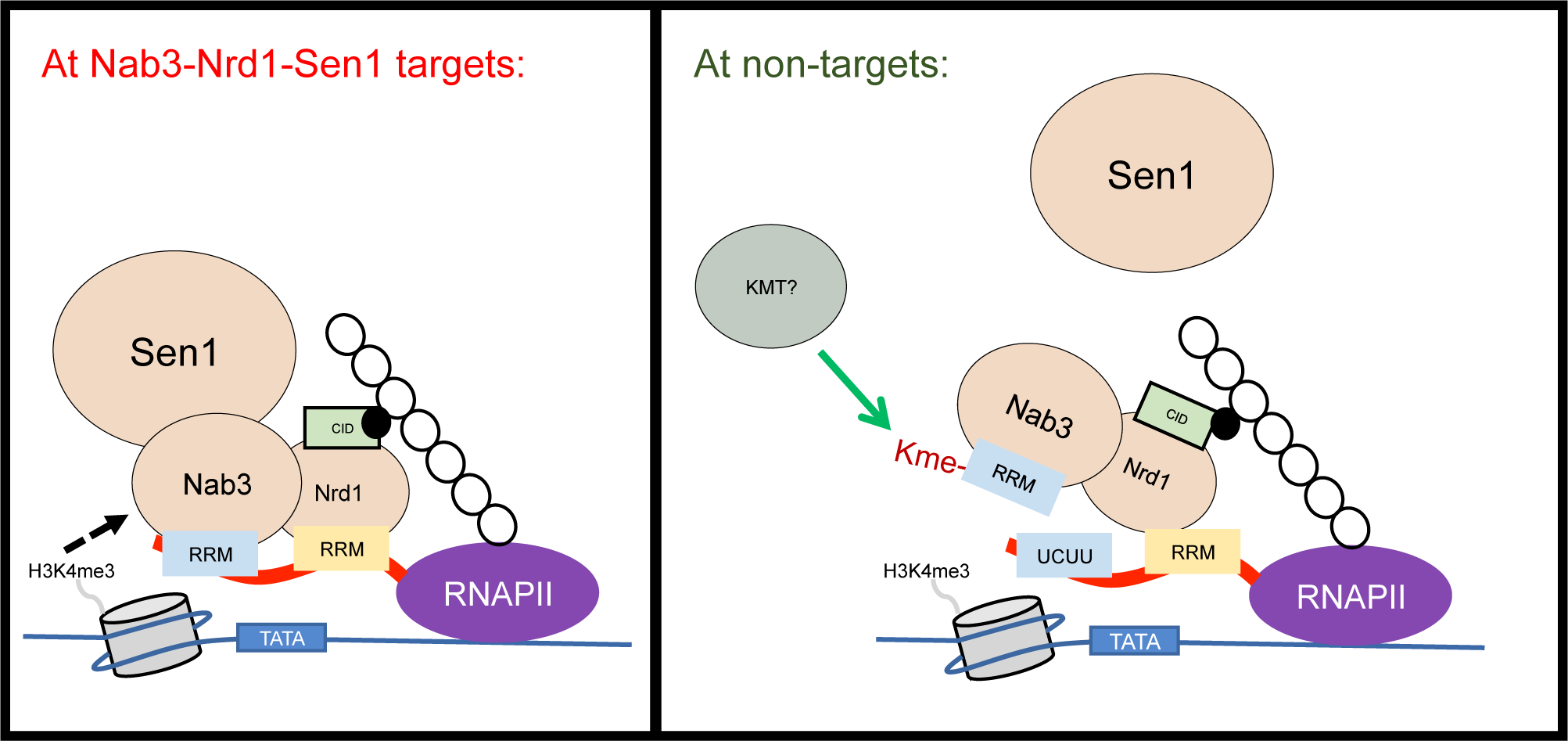
Hypothesized functional significance of Nab3-K363 methylation. Nab3 and Nrd1 integrate numerous molecular inputs to target short noncoding RNAs for termination (left). Methylation of Nab3-K363 may attenuate its RNA binding affinity, leading to regional down-regulation of NNS termination (right).

Recent research has shown that numerous non-histone proteins undergo dynamic lysine methylation and that these modification states have important regulatory functions for the respective proteins (55-57). These studies suggest a widespread role for lysine methylation in regulating protein function well beyond modulating chromatin via histone methylation. For example, the tumor suppressor p53 is methylated on multiple lysine residues and these individual modifications regulate p53 function through a surprisingly diverse array of mechanisms (58-60). Further, the catalytic subunit of DNA-dependent protein kinase (DNA-PK), an important regulator of DNA damage repair, is methylated on multiple lysine residues, which dictates DNA-PK’s recruitment to methyl-binding partners and its ability to effectively repair damaged DNA (56). More recently, Tubulin, Elongin A, and JARID2 have been identified as targets of the human histone H3-K36 and H3-K27 methyltranseferases SETD2 and PRC2, respectively (61-63). Indeed, a growing number of methyltranseferases have been found to modify both histone and non-histone proteins, and several have been shown to be specific for non-histone proteins. As such, it is not surprising that the lysine methylation of non-histone proteins has emerged as an important PTM with wide-ranging impact over a growing number of cellular processes and disease states.

Of the nine NNS lysine methylations we identified, only Nab3-K363 exhibited an obvious functional role. Of course, our studies are not sufficient to rule out any functional roles for the eight additional methylation sites, and more detailed interrogation will be needed to address this, including the identification of the methyltransferases that deposit their methylations. Our studies of Nab3-K363 have homed in on RNA binding as the critical function for this residue, and structural studies have revealed that Nab3-K363 forms a hydrogen bond with the RNA backbone of a target RNA (Figure 4A) (37,38). The RNA binding properties of wildtype, Nab3-K363A, and Nab3-K363R reveal that the precise biochemical properties of Nab3-K363 are crucial for Nab3 RNA binding and NNS function. While this is not surprising given that Nab3-K363 forms a hydrogen bond with the RNA backbone, they do suggest that the methylation of Nab3-K363 may alter its binding, perhaps through charge shielding or altering the hydrophobic character and/or size of the modified lysine (Figure 4A and 5). To test this hypothesis, it will be necessary to identify the methyltransferase(s) that deposit Nab3-K363 mono and trimethylation to facilate *in vitro* methylation of Nab3-K363 and biochemical interrogation using RNA binding assays.

To our knowledge, the findings described here represent the first investigation of any non-histone lysine-methylation of a chromatin-associated protein in yeast. Given the abundance of known or suspected methyltransferases that are associated with yeast chromatin, it seems likely that our findings represent a rule, not an exception. Specifically, a prediction of our findings is that many, or perhaps most chromatin-associated proteins possess lysine methylations, and that at least some of these may be of regulatory significance. Indeed, although numerous lysine methyltransferases have been highly characterized with respect to their ability to control methylation at specific histone residues, the known targets of these enzymes remains largely restricted to their respective histone substrates. Both in yeast as well as in other organisms, whether the regulatory impact of these methyltransferases is explained solely through their histone substrates remains an open question. It will be interesting to identify the methyltransferases responsible for the NNS methylations we describe here to test and extend our model. More broadly, a comprehensive and systematic description of the chromatin “methyl-ome” is an attractive line of investigation.

## DATA AVAILABILITY

All data is available upon request.

## SUPPLEMENTARY DATA

Supplementary Data are available at NAR online.

## ACKNOWLEDGEMENTS

We are grateful to Dr. David A. Brow (University of Wisconsin) for *SEN1* plasmids and the Nrd1 and Sen1 antibodies. We thank Dr. Trevor F. Moraes (University of Toronto) for providing the pET28A expression vector used in this study. Dr. Maurice Swanson at the University of Florida kindly provided the Nab3 antibody. Heterozygous *nrd1Δ*/NRD1, *nab3Δ/NAB3, sen1Δ/SEN1* strains were generously provided by Dr. Charlie Boone and Dr. Brenda Andrews.

## TABLE AND FIGURES LEGENDS

**Figure S1.**
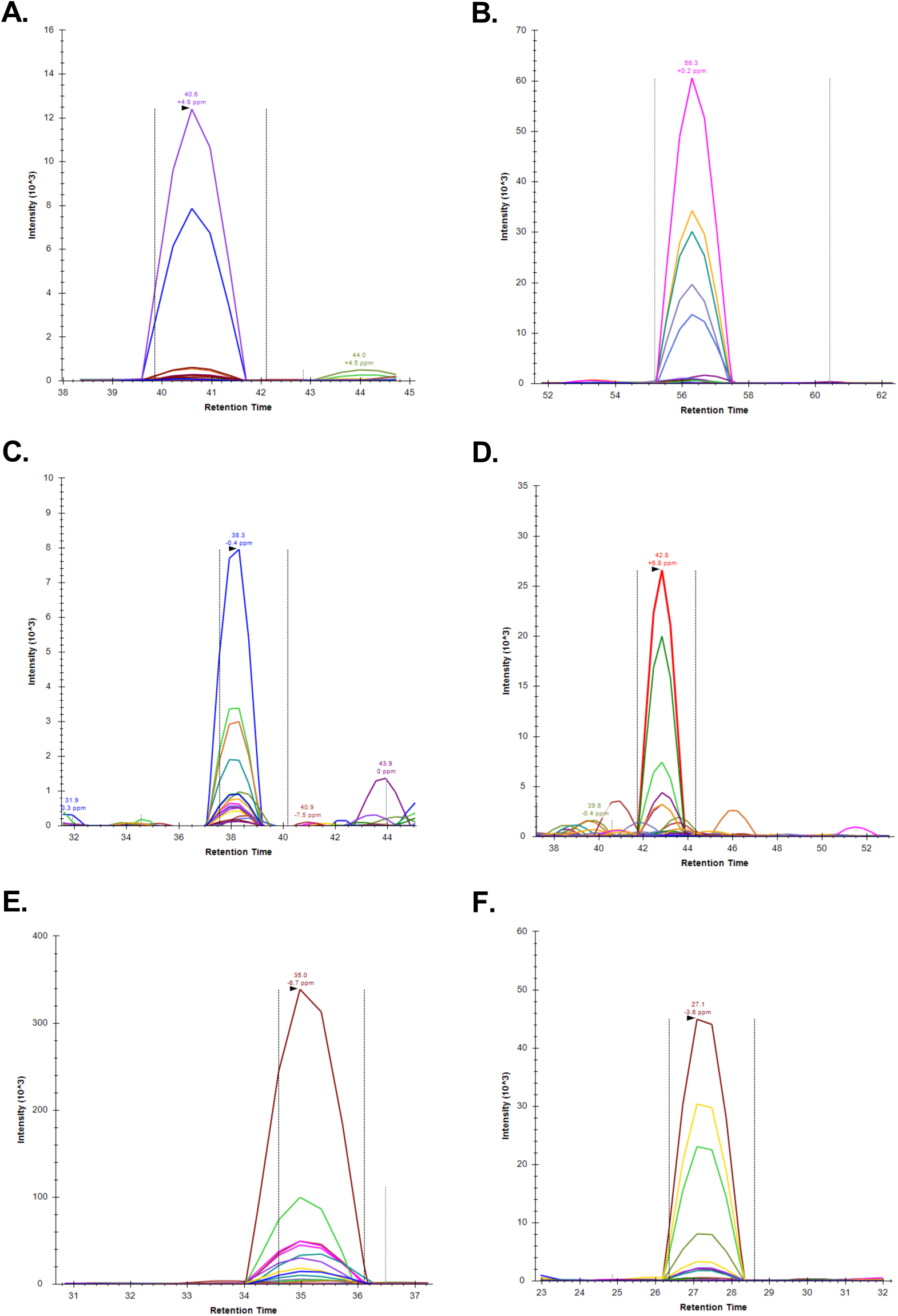
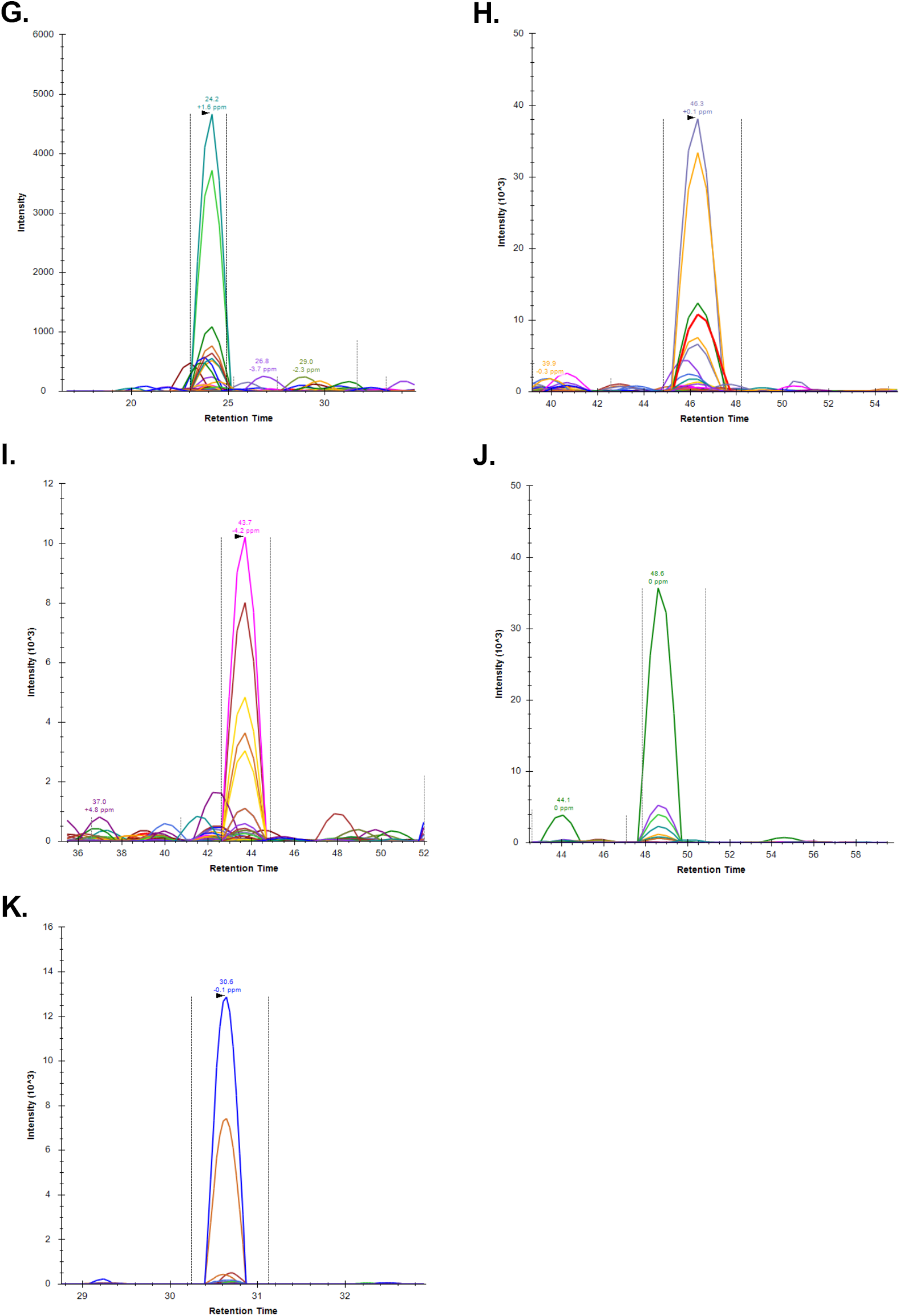
SRM-MS based identification of novel Nab3, and Sen1 lysine methylation events. Transition ions for (A) Nab3(K73me1), (B) Nab3(K213me1), (C) Nab3(K363me3), (D) Nab3(K393me2), (E) Sen1(K19me1), (F) Sen1(K19me2), (G) Sen1(K21me1), and (H) Sen1(K21me2) methylations are shown. Chromatographs were compiled by Skyline v.2.5.0.5675 software.

**Figure S2.**
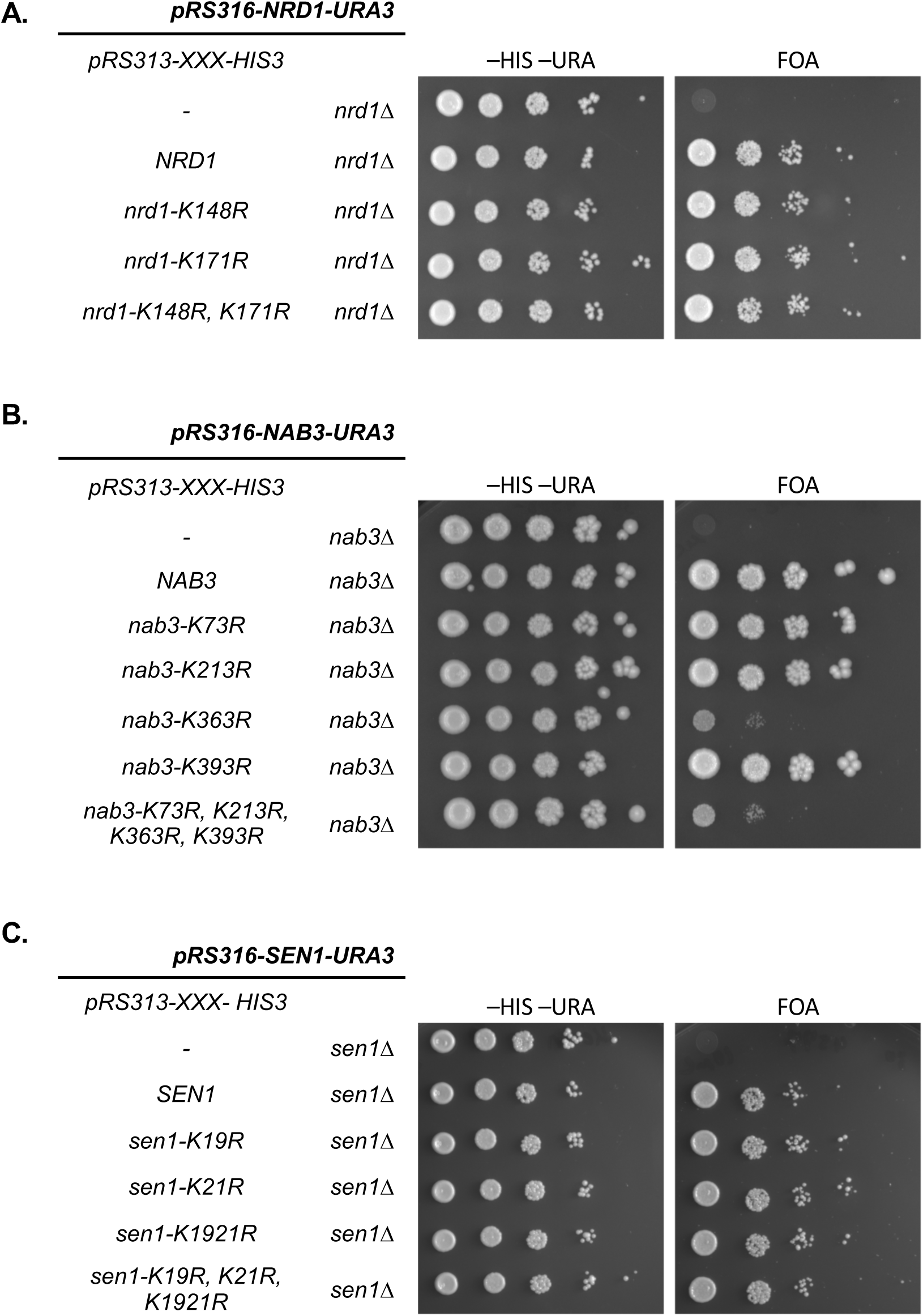
Systematic mutation of methylated lysine residues to arginine. Yeast strains possessing the indicated plasmids were serially diluted ten-fold, spotted onto agar plates containing -HIS -URA dropout media and also on synthetic complete media supplemented with 5FOA. The “-” symbol represents pRS313-HIS3 (empty vector). All strains were grown at 30°C. Systematic mutation of (A) Nab3, (B) Nrd1, (C) Sen1 methylated lysine residues to arginine.

**Figure S3.**
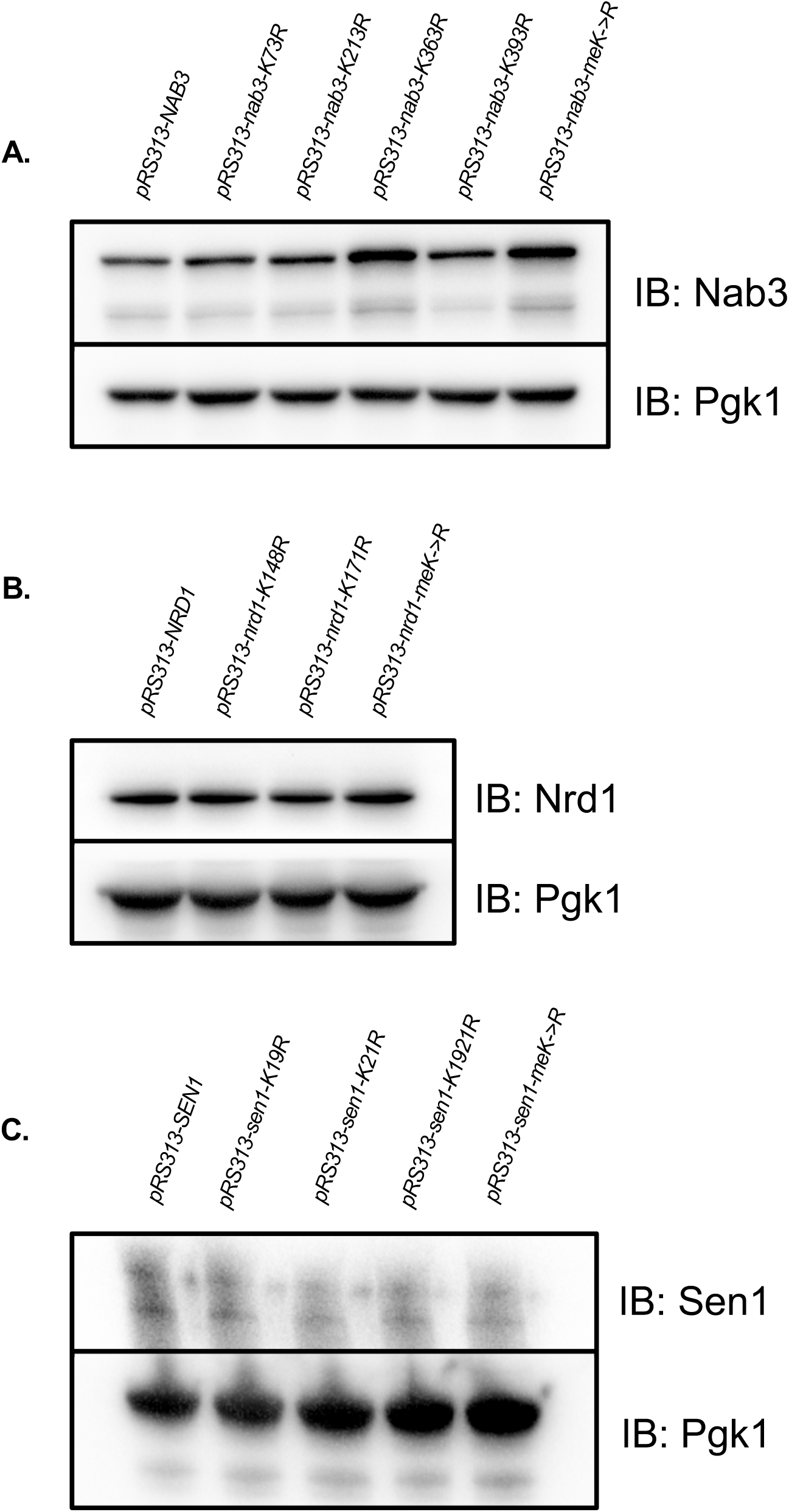
Nrd1, Nab3, Sen1 K to R mutants all express stable proteins. Immunoblot analysis of (A) Nab3, (B) Nrd1, and (C) Sen1 protein levels in wildtype and respective lysine to arginine substitution mutants. meK-->R represents mutants with all methylated lysine residues mutated to arginine.

**Figure S4.**
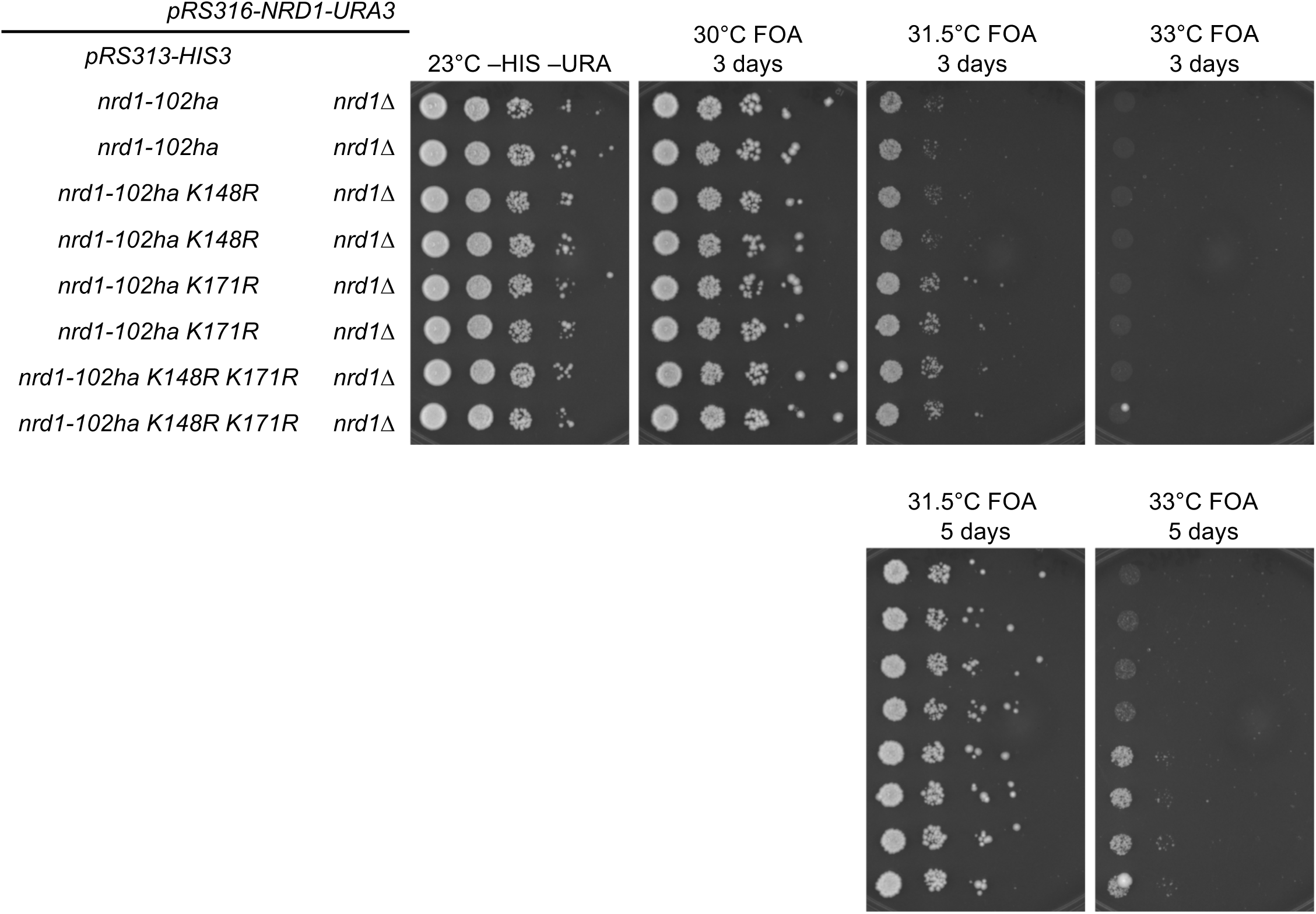
*nrd1-102ha* mutants are weakly suppressed by a K171 to arginine substitution. Yeast strains possessing the indicated plasmids were serially diluted ten-fold, spotted onto agar plates containing -HIS -URA dropout media and also on synthetic complete media supplemented with 5FOA. Growth of two independent isolates of each genotype is shown. Plates at 23°C -HIS-URA and 30°C FOA are shown after 3 days of growth, while plates at 31.5°C, and 33°C FOA are shown after 3 days and 5 days of growth.

**Figure S5.**
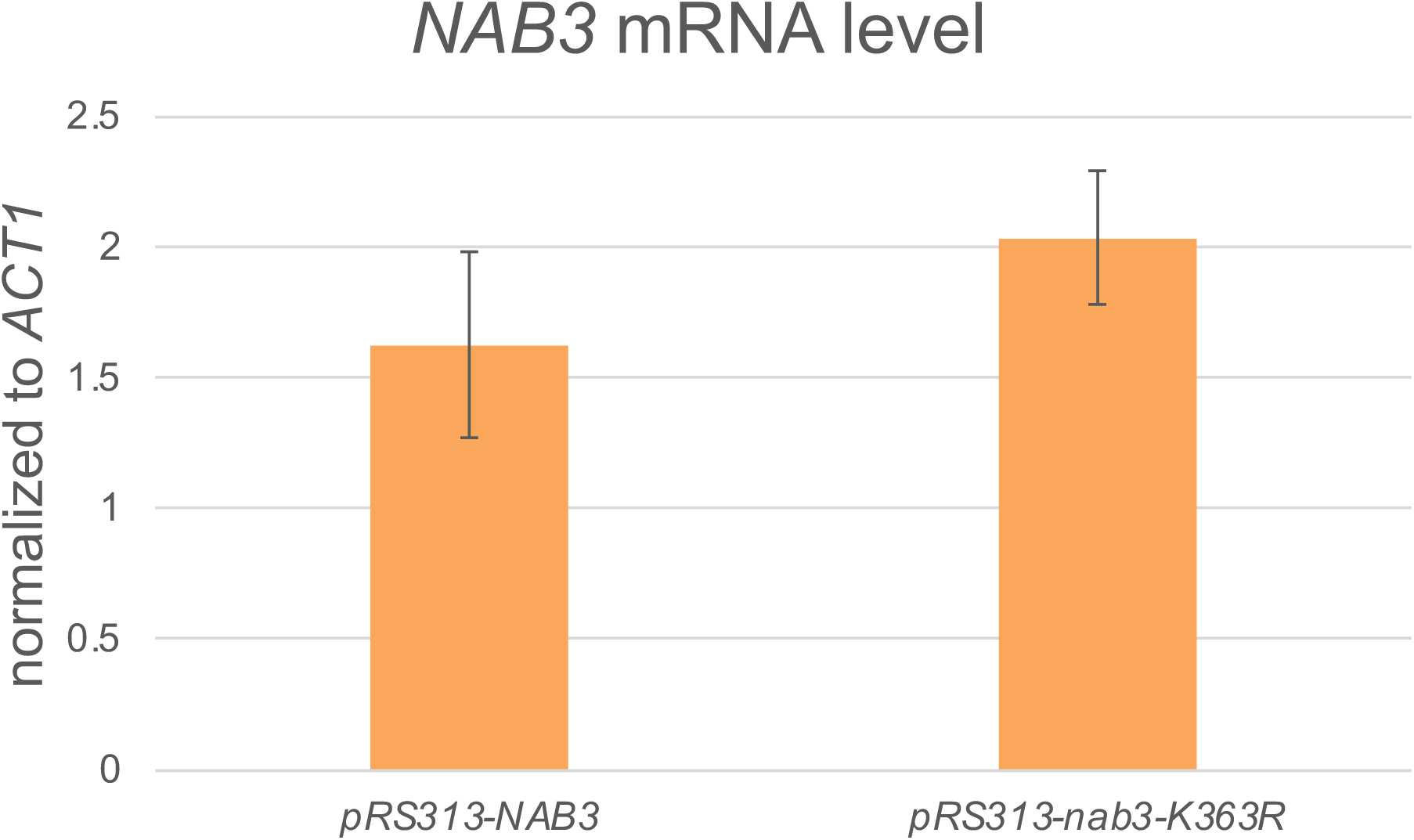
Steady state level of the *NAB3* mRNAs. *NAB3* mRNA level from *NAB3* and *nab3-K363R* cells was analyzed by qRT-PCR. The average of three biological replicates is shown. Error bars represent standard deviation. Significance is calculated by a two-tailed student’s t-test p>0.05, n=3.

**Figure S6.**
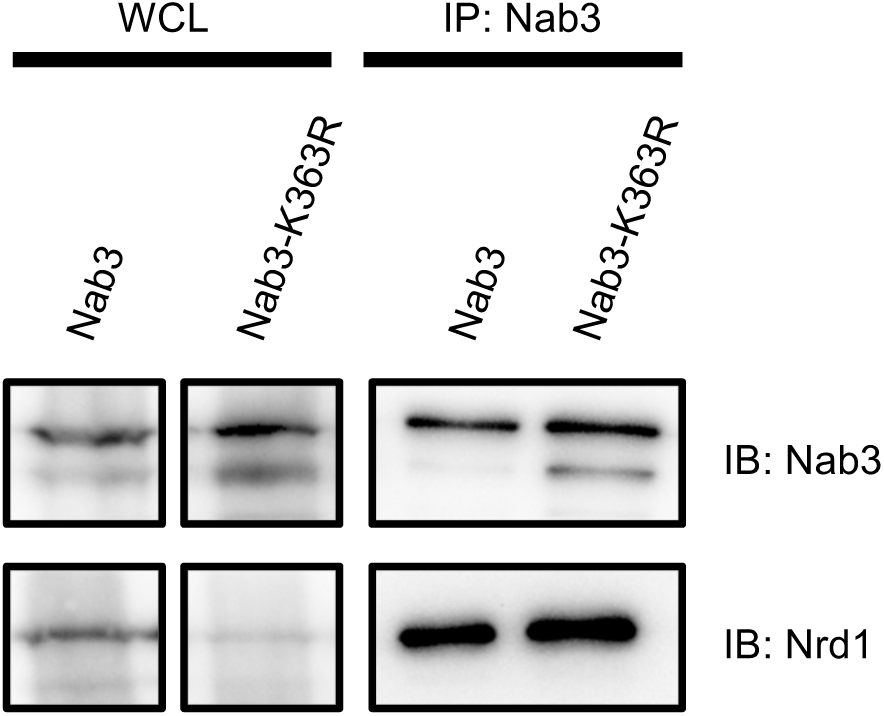
Nab3 immunoprecipitation in wildtype and *nab3-K363R* mutants. Total cell lysates (WCL) were immunoprecipitated with a monoclonal Nab3 antibody (IP: Nab3) followed by immunoblot analysis with anti-Nab3 and anti-Nrd1 antibodies.

**Figure S7.**
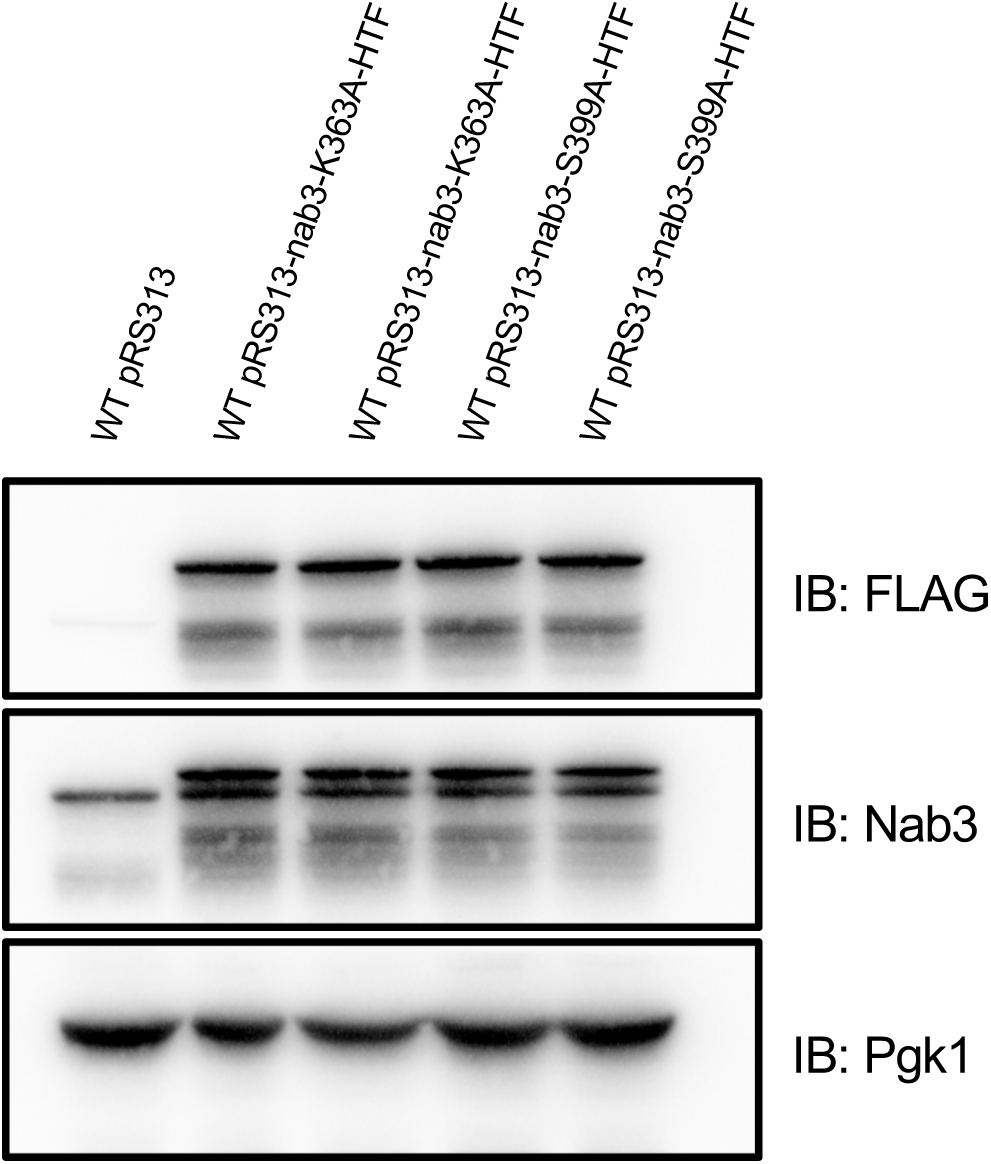
Nab3-K363A and Nab3-S399A plasmid constructs express stable proteins. Immunoblot analysis of wildtype cells expressing *pRS313* (empty vector), *pRS313-nab3-K363A-HTF*, and *pRS313-nab3-S399A-HTF*. Nab3-K363A-HTF (His_6_-TEV-FLAG) and Nab3-S399A-HTF protein were detected by an anti-FLAG antibody as well as the anti-Nab3 antibody. On the Nab3 blot, the top band likely represents the Nab3-HTF construct while the bottom band represents endogenous Nab3. Pgk1 protein level was detected using the anti-Pgk1 antibody and used as a loading control.

**Figure S8.**
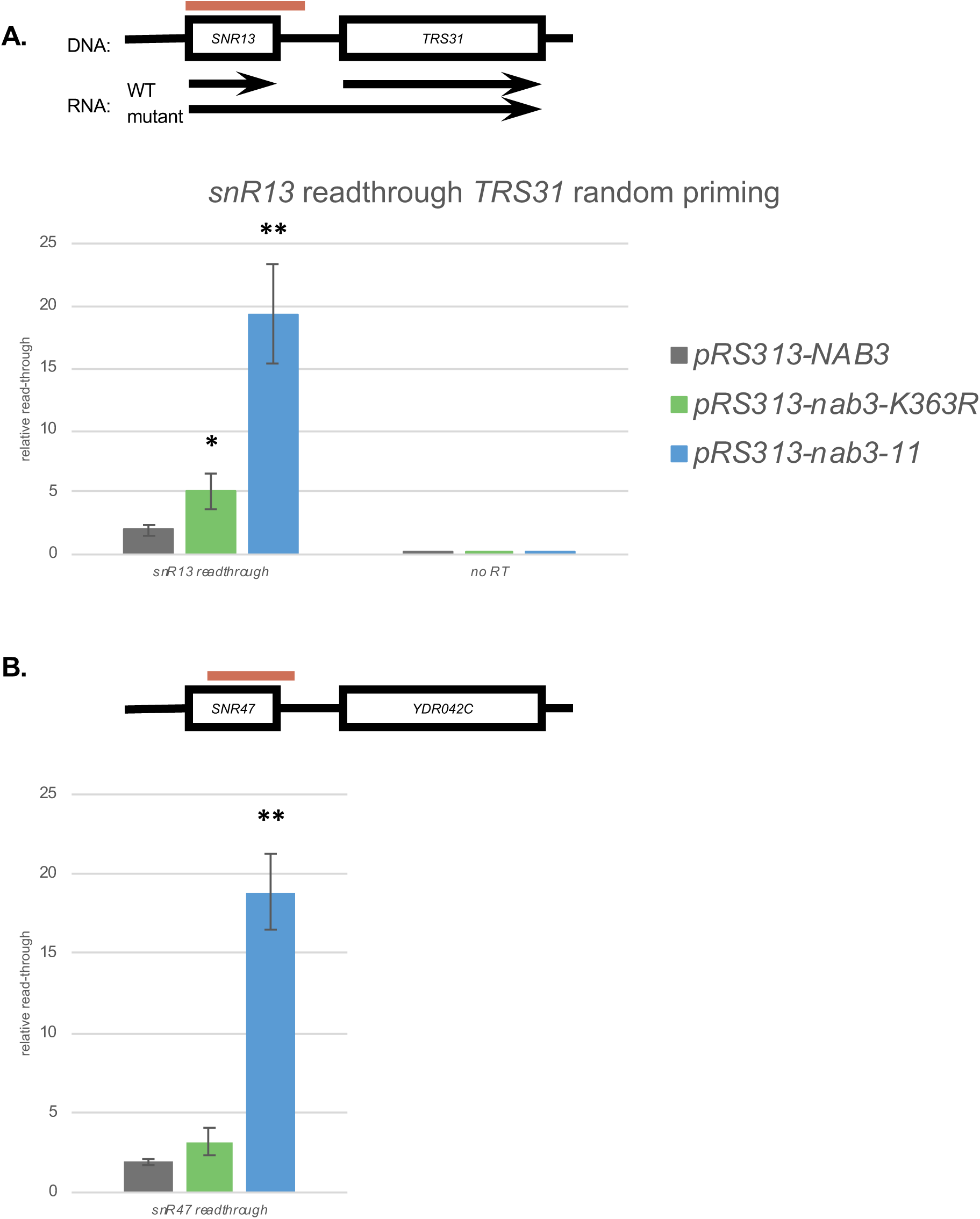
qRT-PCR analysis of *NAB3, nab3-K363R*, and *nab3-11* cells. (A) *TRS31* random priming matches the results seen for gene specific priming. Total RNA from *NAB3, nab3-K363R,* and *nab3-11* cells was processed into cDNA by random nonamers instead of a *TRS31* gene specific primer. The presence of *SNR13* read-through transcription was analyzed by qPCR. (B) The level of read-through transcription at *SNR47*. Error bars represent standard deviation of three biological replicates. Significance between each mutant and the WT control is calculated by a two-tailed student’s t-test and denoted by * p<0.05, ** p<0.01, n=3. No reverse transcription control (No RT, n=1).

**Table S1.**
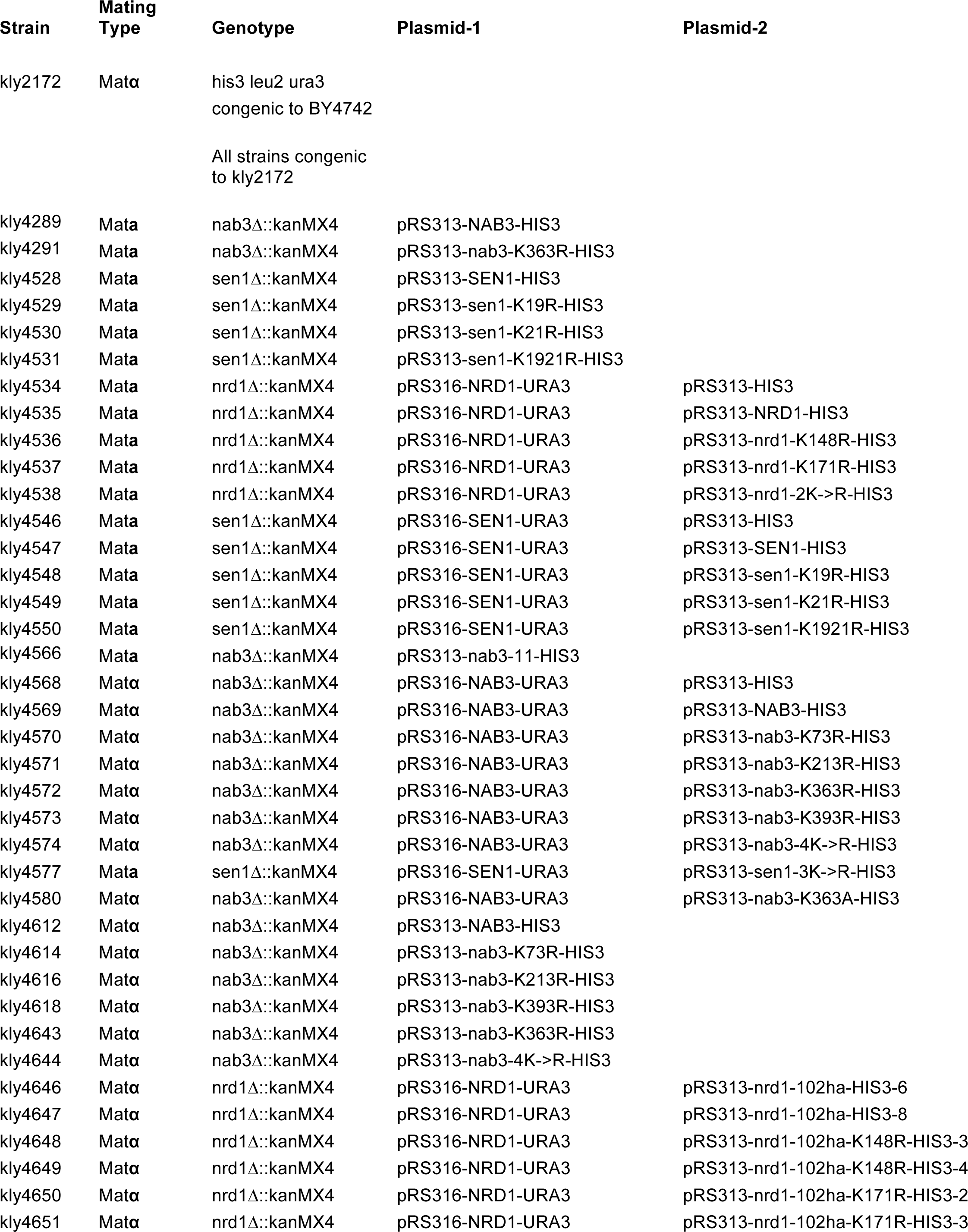
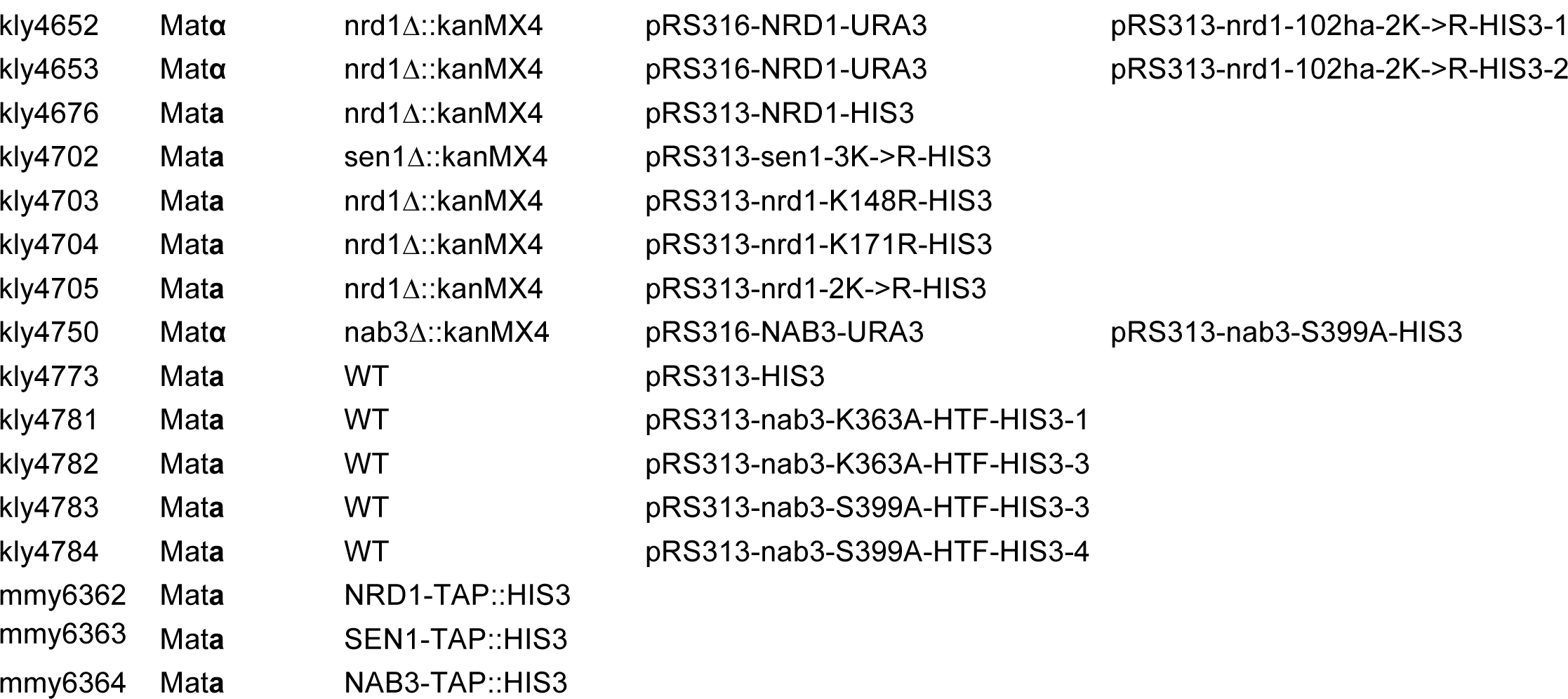

**Table S2.**
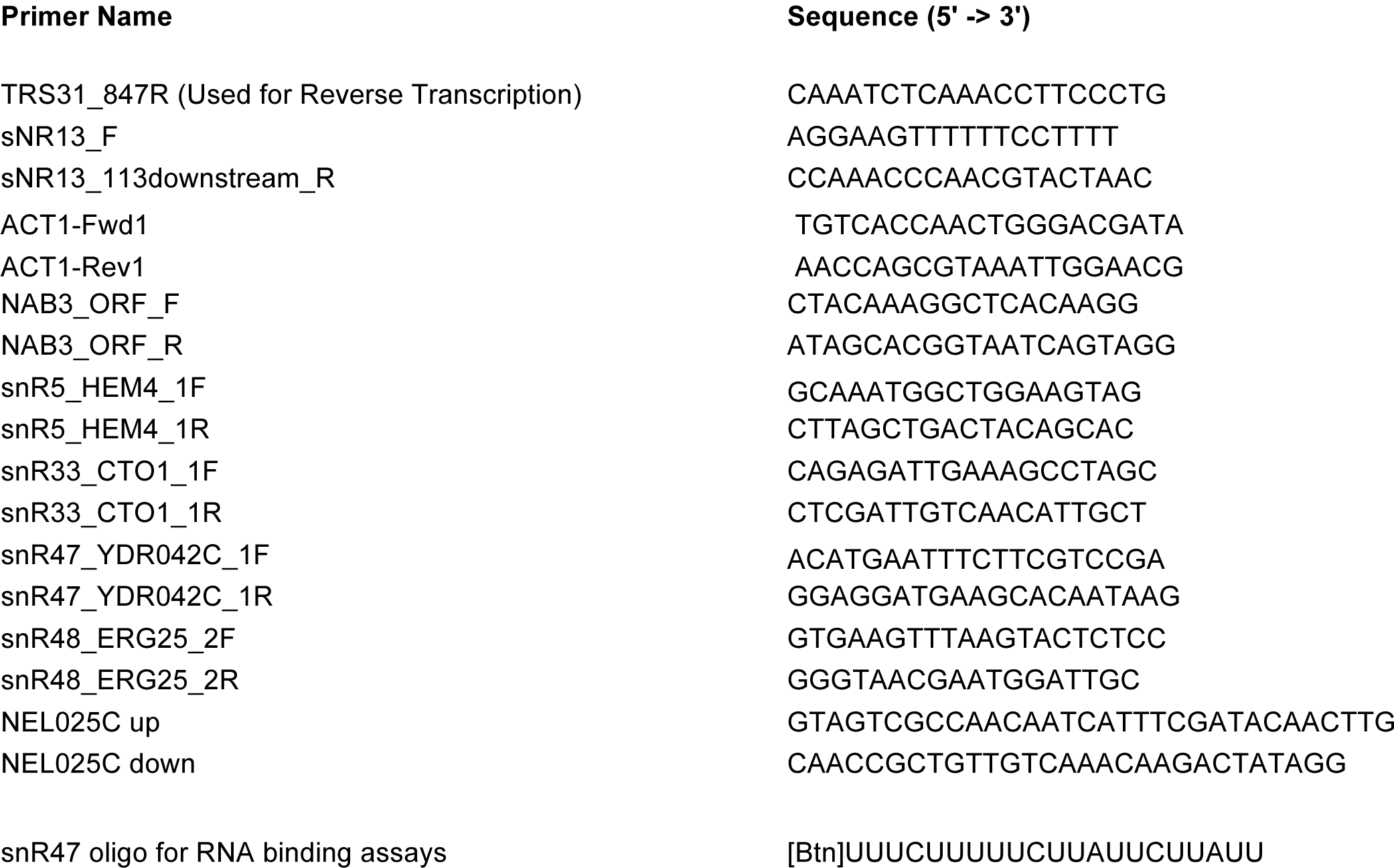

